# Activation of Vasopressin Receptor 1A by Vasopressin Enhances Myometrial Smooth Muscle Cell Excitability by Inhibiting the Potassium Channel SLO2.1

**DOI:** 10.64898/2026.08.06.743275

**Authors:** Juan Ferreira, Lindsey N. Kent, Ana Gonzalez-Cota, Nikita Peramsetty, Grace C. Whitter, Ethan Li, Sariela Spivak, Xiaofeng Ma, Sarah K. England, Celia M. Santi

## Abstract

Arginine vasopressin (AVP) increases excitability of myometrial smooth muscle cells (MSMCs) through Gαq-coupled AVP receptors. Although excitability requires membrane depolarization, the mechanisms linking AVP receptor activation to membrane depolarization and Ca²⁺ signaling are incompletely understood. Here, we show that AVPR1 is the predominant AVP receptor in primary MSMCs. In *Xenopus* oocytes, AVP signals through AVPR1 to inhibit SLO2.1-mediated potassium currents, reducing current amplitude to approximately 60% of control currents. Consistent with suppression of a hyperpolarizing conductance, AVP depolarized a myometrial cell line (hTERT-HM) and increased intracellular Ca²⁺ signaling. Analysis of Ca²⁺ dynamics revealed that the initial Ca²⁺ peak was largely preserved under conditions limiting extracellular Ca²⁺ entry, consistent with intracellular store release. Conversely, the oscillatory phase depended on extracellular Ca²⁺ influx and was reduced by SLO2.1 knockdown. Together, these findings support a model in which AVP preferentially signals through AVPR1A to inhibit SLO2.1, depolarize myometrial cells, enhance VDCC-dependent Ca²⁺ entry, and promote excitability, enhancing conditions for uterine contraction.

## Introduction

Pregnancy requires coordinated adaptations of the reproductive and endocrine systems to support fetal development and prepare the uterus for parturition. Two key signaling systems that undergo pregnancy-associated changes are the vasopressinergic and oxytocinergic systems (Fenelon, Poulain and Theodosis, 1993; Lin *et al*., 1996; Landgraf and Neumann, 2004; Koshimizu *et al*., 2012; Szczepanska-Sadowska *et al*., 2017; Lozić *et al*., 2018). Arginine vasopressin (AVP) and oxytocin (OXT) are synthesized predominantly by hypothalamic neurons and released into the circulation from the posterior pituitary in response to physiological stimuli (Sofroniew, 1983; Tribollet and Barberis, 1996; Dai *et al*., 1998). In addition, AVP and OXT are produced by the uterus and placenta, where they contribute locally to regulation of pregnancy-related processes (Stegner *et al*., 1984; Oosterbaan, Swaab and Boer, 1985; Oosterbaan and Swaab, 1989; Irion, Mack and Clark, 1990; Patient *et al*., 1999; Downing *et al*., 2016) AVP and OXT are closely related nonapeptide hormones that differ by only two amino acids and are thought to have evolved from a common ancestral peptide gene (Carter, 2017). Their receptors likewise belong to the same subfamily of G protein-coupled receptors and share substantial structural and functional similarities (Tribollet and Barberis, 1996; Landgraf and Neumann, 2004; Koshimizu *et al*., 2012). Although each peptide preferentially activates its cognate receptors, their structural similarity and close evolutionary relationship permit some degree of cross-receptor activation (MAGGI *et al*., 1990; Song *et al*., 2014; Song and Albers, 2018; Elsaafien *et al*., 2026).

Although AVP is best known for its roles in fluid homeostasis and cardiovascular regulation, several features make its actions relevant to uterine smooth muscle (myometrium). First, pregnancy resets the osmotic threshold for AVP release and circulating AVP concentrations increase despite enhanced degradation by placental vasopressinase (Davison *et al*., 1988, 1989, 1993; Lindheimer, Barron and Davison, 1989; Lindheimer and Davison, 1995; Tkachenko, Shchekochikhin and W. Schrier, 2014). Second, AVP release is regulated not only by osmotic and hemodynamic stimuli but also by pain, stress, hypoxia, and bleeding, all of which are relevant during pregnancy and labor (KENDLER, WEITZMAN and FISHER, 1978; Suzuki *et al*., 2009; Sivukhina *et al*., 2010; Szczepanska-Sadowska *et al*., 2017). Third, increased vasopressinergic signaling has been associated with adverse pregnancy outcomes, including preeclampsia (Santillan *et al*., 2014; Scroggins *et al*., 2018). Fourth, AVP stimulates the excitability and contraction of both vascular and uterine smooth muscle (Fuchs *et al*., 1992; Lindheimer and Davison, 1995; MASTORAKOS and ILIAS, 2003; Tkachenko, Shchekochikhin and W. Schrier, 2014). Although these observations indicate that AVP signaling contributes to uterine physiology, particularly during pregnancy and labor, the mechanisms underlying the effects of AVP in human myometrial cells are unclear.

Given the similarities between AVP and OXT, the two hormones may regulate the myometrium through similar mechanisms. In mouse and human models, OXT stimulates myometrial excitability/contractility by activating the Gαq/11-coupled oxytocin receptor (OTR), which stimulates phospholipase C (PLC) to hydrolyze phosphatidylinositol 4,5-bisphosphate to generate inositol 1,4,5-trisphosphate (IP₃) and diacylglycerol (DAG) (Exton, 1996; Katan, 2005; Putney and Tomita, 2012; Lyon *et al*., 2023; Ubeysinghe *et al*., 2023). This leads to two branches of signaling, both of which lead to cytosolic calcium (Ca^2+^) increase. In the canonical branch, IP₃ binds to IP₃ receptors on intracellular Ca²⁺ stores, triggering Ca²⁺ release into the cytosol. In the non-canonical branch, DAG activates protein kinase C (PKC), which leads to inhibition of the sodium-activated potassium (K⁺) channel SLO2.1. The resulting reduction in outward K⁺ conductance depolarizes the myometrial cell membrane, increases the open probability of voltage-dependent Ca²⁺ channels, and promotes extracellular Ca²⁺ entry (Ferreira *et al*., 2019, 2021, 2025). The increased cytosolic Ca²⁺ from both branches activates Ca²⁺/calmodulin–myosin light-chain kinase to promote actin–myosin interactions and myometrial contractility.

Unlike OXT, which acts primarily through OTR, AVP signals through three G protein-coupled receptor subtypes: AVPR1A, AVPR1B, and AVPR2. AVPR1A is the most relevant in pregnancy because it is expressed in the myometrium and couples to Gq/11 and activates PLC-dependent production of IP₃ and DAG (Tribollet and Barberis, 1996; Thibonnier *et al*., 2002; Malik *et al*., 2022; Fang *et al*., 2024). Although AVPR1B also couples to Gq/11, it is expressed predominantly in the anterior pituitary and selected regions of the central nervous system. AVPR2 couples primarily to Gs proteins, activates cAMP/PKA signaling, and is expressed mainly in the kidney (Góźdź *et al*., 2002; Mutig *et al*., 2007; El-Werfali *et al*., 2015). Given the shared signaling architecture of AVPR1A and OTR, we hypothesized that AVP acts through AVPR1A in myometrial cells to increase cytosolic Ca^2+^ via two pathways. In one, AVP would promote IP₃-dependent signaling to mobilize Ca²⁺ from intracellular stores. In the second, AVP would activate DAG/PKC signaling to inhibit SLO2.1-mediated K⁺ conductance, depolarize the resting membrane potential, and enhance extracellular Ca²⁺ entry through voltage-dependent Ca²⁺ channels.

To test this hypothesis, we examined AVPR1A expression in primary myometrial smooth muscle cells (MSMCs) isolated from women at term, both before labor and during labor. We then examined the effects of AVP on SLO2.1 currents, the resting membrane potential, and intracellular Ca²⁺ signaling in immortalized human myometrial (hTERT-HM) cells. Finally, we used targeted knockdown of SLO2.1 and targeted knockdown or knockout of AVPR1A and OTR to define the receptor and ion-channel specificity of the effects of AVP stimulation.

## Methods

This study did not generate new unique reagents.

### Ethical approval and acquisition of human samples

This study was approved by the Washington University in St. Louis Institutional Review Board (protocol no. 201108143) and conformed to the Declaration of Helsinki except for registration in a database. We obtained signed written consent from each patient. Human tissue samples (0.1-1.0 cm^2^) from the lower uterine segment were obtained from term non-laboring (TNL) and term laboring (TL) women (>37 weeks of gestation) during elective Cesarean section under spinal anesthesia. Samples were stored at 4°C in phosphate-buffered saline (PBS) and processed for MSMC isolation within 60 min of acquisition.

### Cell culture

HEK293T and hTERT cells where cultured in Dulbecco’s Modified Eagle Medium (DMEM)/Ham’s F12 media without phenol red supplemented with 10% fetal bovine serum and 25 μg/ml gentamicin (all from Sigma, St. Louis, MO). Cells were cultured in a standard humidified cell culture incubator at 37°C with 5% CO_2_. The hTERT cell line was a kind gift from Dr. Jennifer Condon at Wayne State University (Condon *et al*., 2002).

### Isolation of primary human MSMCs

Primary MSMCs were generated as previously reported (Malik *et al*., 2022). Briefly, myometrial samples were disassociated in Collagenase IA and Collagenase XI (1 mg/ml each, Millipore Sigma) at 37°C for at least 45 min. Collagenases were inactivated by adding DMEM/Ham’s F12 media containing 10% fetal bovine serum and 25 μg/ml gentamicin, and cells were passed through a 100 μm cell strainer. After centrifugation, the pellet was re-suspended in DMEM/Ham’s F12 media supplemented with 5% Smooth Muscle Cell Growth Medium 2 (PromoCell) and 25 μg/ml gentamicin and plated.

### Quantitative Real-time PCR

Total RNA was isolated from hTERT cells and hMSMCs by using an RNeasy Plus Kit (Qiagen, Germantown, MD). Complementary DNA was made from 0.5–1 µg of RNA with iScript RT Supermix (Bio-Rad, Hercules, CA) and amplified with iQ SYBR Green Supermix (Bio-Rad). Samples were amplified in a Bio-Rad CFX Connect Real-Time PCR Detection System with the following temperature specifications: 95°C for 3 min (x1), then 95 °C for 10 s and 60°C for 30s (40x). Abundance of each transcript was normalized to the reference gene Topoisomerase I (*TOP1*) by the ΔΔCt method.

### Electrophysiology

Cells were starved in serum-free DMEM:F12 for at least two hours before experiments. For all experiments, pipettes were pulled from borosilicate glass from Warner Instruments. For whole-cell recording, pipettes with a resistance of 0.8 to 1.8 megaohms and symmetrical K^+^ were used. External solution was (in mM): 160 KCl, 80 NaCl, 2 MgCl_2_, 10 HEPES, and 5 TEA, pH adjusted to 7.4 with NaOH. For the 0 mM Na^+^ solution, Na^+^ was replaced with 80 mM CholineCl, and pH was adjusted with KOH; the concentration of external K^+^ varied from 4.5 to 5.5 mM. The pipettes were filled with (in mM): 160 KCl, 80 cholineCl or 80 NaCl, 10 HEPES, 0.6 free Mg^2+^, and either 0 or 100 nM free Ca^2+^ solutions with 1 mM EGTA. Variations in the solutions are indicated in the figures. During electrophysiological experiments, the cells and the intracellular side of the membrane were perfused continuously. Traces were acquired with an Axopatch 200B (Molecular Devices), digitized at 10 kHz for whole-cell or macro-patch recordings or at 100 kHz for single-channel recordings. Recordings were filtered at 2 kHz, and pClamp 10.6 (Molecular Devices) and SigmaPlot 15 (Jandel Scientific) were used to analyze the data.

### Two-electrode voltage clamp experiments in *Xenopus* oocytes

Oocytes were harvested from female *Xenopus laevis* as described by Yuan et al. (2000)(Yuan *et al*., 2000). Defolliculated oocytes were injected with 46 and 92 ng of cRNA with a nano injector (Drummond Scientific, Broomall, PA, USA). Injected oocytes were kept at 18°C in ND96 medium containing (in mm): 96 NaCl, 2 KCl, 1.8 CaCl_2_, 1 MgCl_2_ and 5 Hepes, pH 7.5 (with NaOH). Two-electrode voltage clamp experiments were performed 3–5 days after oocyte injection, as described previously (Wei et al. 1994). Whole-cell current recordings from *Xenopus* oocytes were performed while the oocytes were being perfused with ND96 + 2 μm 4,4′-diisothiocyanatostilbene-2,2′-disulphonic acid (catalogue no. D3514; Sigma-Aldrich) to block endogenous chloride currents. All records were obtained with the voltage protocols specified. Data were acquired with a Digidata 1440 (Molecular Devices). Oxytocin (OXT, catalogue no. 1910; Tocris Bio-Techne Corporation, Minneapolis, MN, USA), and arginine vasopressin (AVP, CAS number 113-79-1; Sigma-Aldrich) were applied to the recording chamber by continuous perfusion.

### Determination of membrane potential by flow cytometry

hTERT-HM cells were centrifuged at 300 g for five minutes. Cells were resuspended in modified Ringer solution containing (in mM): 135 NaCl, 10 HEPES, 5 Glucose, 5 KCl, and 2 CaCl_2_; pH 7.4. Before recording, 0.02 mg/mL Hoechst and 150 nM DiSC_3_(5) were added to 500 μL of cell suspension, and data were recorded as individual cellular events. Side scatter area (SSC-A) and forward scatter area (FSC-A) fluorescence data were collected from 100,000 events per recording. Thresholds for FSC-A and SSC-A were set to exclude signals from cellular debris. Doublets, aggregates, and cell debris were excluded from the analysis based on a dual parameter dot plot in which pulse signal (signal high; SSC-H; y-axis) versus signal area (SSC-A; x-axis) was displayed. Living Hoechst-negative cells were selected by using the filter Pacific Blue (emission 450±25 nm), and DiSC3(5)-positive cells were detected with the filter for allophycocyanine (emission 660±10 nm).

To measure the effect on membrane potential, OXT, phorbol 12-myristate 13-acetate (PMA), or AVP were added to 500 μL of cell suspensions. Valinomycin (1 μM) (Sigma, St. Louis, MO), which hyperpolarizes the cell to the equilibrium potential for K^+^, was used to establish a reference across different experiments. Normalization was performed according to the forumula (F_X_-F_Ref_)/(F_Valino_-F_Ref_), where F_Ref_ is the median of the fluorescence of the population in the basal condition, F_X_ is fluorescence after addition of OXT, PMA or AVP, and F_Valino_ is fluorescence after addition of Valinomycin. FlowJo 10.6.1 software was used to analyze data, reported as median values.

### Calcium imaging

hTERT-HM cells were grown on glass coverslips with DMEM/Ham’s F12 media containing 10% fetal bovine serum. Cells were pre-incubated with 2 μM Fluo-4 AM and 0.05-0.1% Pluronic Acid F-127 in Opti-Mem for 60-90 min. To allow the dye to equilibrate in the cells, the cells were removed from the loading solutions and placed in Ringer solution for 10 to 20 minutes. The various solutions were applied with a perfusion system with an estimated exchange time of 1.5 s. Recordings started 2–5 min before addition of the first test solution. Ionomycin (5 μM) was added at the end of the recordings as a control stimulus. Calcium signals were recorded with a Leica AF 6000LX system with a Leica DMi8000 inverted microscope and an Andor-Zyla-VCS04494 camera. A halogen lamp was used with a 488 ± 20 nm excitation filter and a 530 ± 20 nm emission filter. A 40X (HC PL FluoTar L 40X/0.70 Dry) or a 20X (N-Plan L 20X/0.35 Dry) air objective were used. Leica LasX2.0.014332 software was used to collect data and control the system. Acquisition parameters were: 120 ms exposure time, 2×2 binning, 512 x 512 pixels resolution, and a voxel size of 1.3 μm for the 20X objective. Whole images were collected every 10 seconds. LAS X, ImageJ, Clampfit 10 (Molecular Devices), and SigmaPlot 15 were used to analyze data. Changes in intracellular Ca^2+^ concentration are presented as (F/F_Iono_) after background subtraction. All imaging experiments were done at room temperature. Cells were counted as responsive if they had changes in fluorescence of at least 5-10% of the ionomycin response.

### Small interfering RNA

hTERT-HM cells and primary MSMCs were transfected with a combination of three 27-mer small interfering RNAs (siRNAs) from OriGene (Rockville, MD, USA) targeting the human SLO2.1 gene (*KCNT2;* catalogue no. SR317593), *AVPR1* (catalogue no. SR300383) or *OXTR*, (catalogue no. SR303320) all at a final concentration of 25 nm each, or with 75 nm scrambled siRNA. Lipofectamine 2000 transfection reagent (catalogue no. 11668-027; Invitrogen) was used in accordance with the manufacturer’s instructions. Cells were also co-transfected with 10 μg mL^−1^ of a vector encoding green fluorescent protein. Experiments were performed 36–72 h after transfection.

### Statistical analysis

Statistical analyses were performed in SigmaPlot (Systat Software Inc.; version 15.0). For comparisons between two independent groups, unpaired Student’s t-tests were used. For case–control designs in which measurements were obtained from the same individual, paired t-tests were used. Comparisons among more than two groups were performed with one-way ANOVA followed by appropriate post-hoc multiple-comparison tests. Data are presented as mean ± SD, and p < 0.05 was considered statistically significant. The specific statistical test used, the number of cells/individuals, and measures of central tendency and dispersion are reported in the figure legends and/or results section. No *a priori* power analysis was performed; therefore, results based on sample sizes <5 should be considered preliminary and p-values interpreted as descriptive.

## Results

### Human myometrial cells express SLO2.1, AVPR1A, AVPR2, and OTR

To determine whether human myometrial smooth muscle cells (MSMCs) are capable of responding directly to AVP, we assessed the expression of vasopressin receptors by quantitative PCR in primary MSMCs from term non-laboring (TNL; n = 4) and term laboring (TL; n = 3) patients, as well as in hTERT-HM cells (n = 8). As expected, transcripts encoding SLO2.1 and OTR were detected in both TL and TNL samples **(Fig. 1A,B).** Transcripts encoding AVPR1A and AVPR2 were detected in all primary MSMC preparations and in hTERT-HM cells **(Fig. 1C,D).** AVPR1A expression was comparable between TL and TNL samples, whereas AVPR2 expression was significantly lower in TL samples than in TNL samples **(Fig. 1C,D).** These findings demonstrate that human myometrial cells express the receptors required to respond directly to AVP.

**Figure 1.**
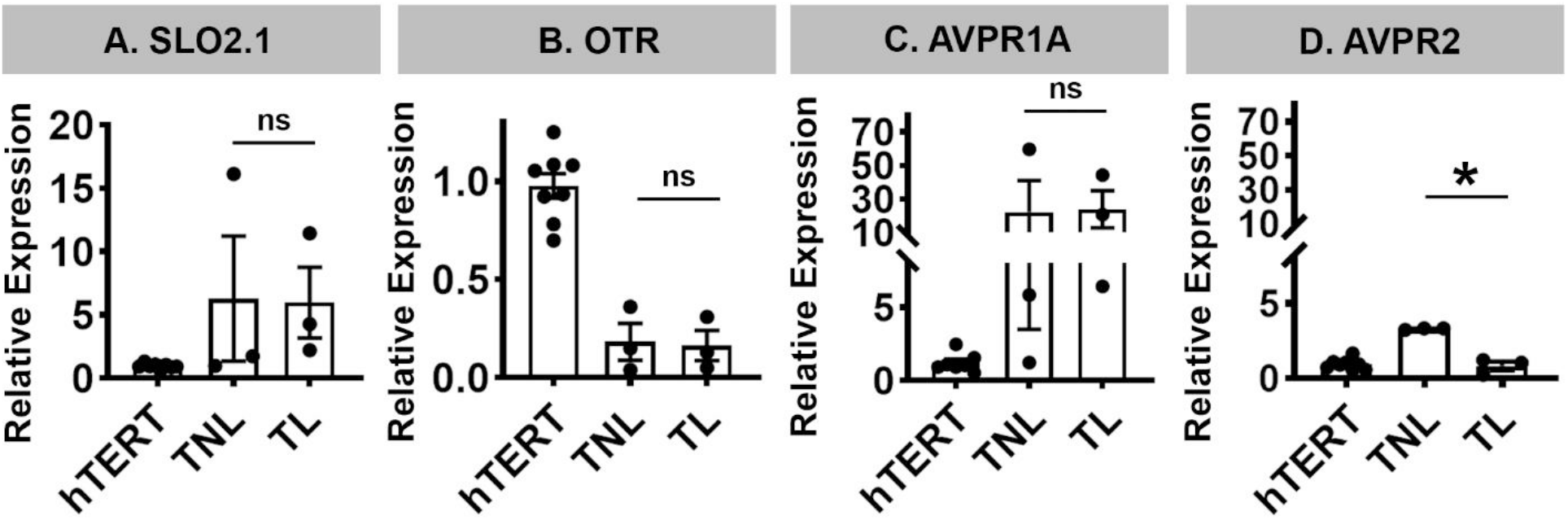
Expression of SLO2.1, OTR, AVPR1A, and AVPR2 in human myometrial cells. Gene expression was assessed by quantitative PCR. **A–B,** Expression of SLO2.1 and OTR in primary myometrial smooth muscle cells from term laboring (TL; *n* = 3), term non-laboring (TNL; *n* = 4) patients, and hTERT-HM Cells (n=8). **C–D,** Expression of AVPR1A and AVPR2 in primary MSMCs from TL and TNL patients and in hTERT-HM cells **(*n* = 8).** Data are presented as mean ± SD. \**P* < 0.05 by unpaired *t* test; ns, not significant.

### Activation of AVPR1A inhibits SLO2.1 currents in a heterologous expression system

To determine whether AVPR1A activation modulates SLO2.1 activity, we co-expressed human SLO2.1 and AVPR1A in *Xenopus* oocytes and measured K⁺ currents via two-electrode voltage clamp **(Fig. 2A).** Application of increasing concentrations of AVP progressively reduced SLO2.1-mediated K⁺ currents, with an apparent IC₅₀ of approximately 15 nM **(Fig. 2A,B)**. In contrast, AVP did not inhibit currents in control oocytes expressing SLO2.1 without AVPR1A **(Fig. 2C)**, indicating that the effect required receptor expression and was not caused by a direct action of AVP on SLO2.1. Oocytes expressing AVPR1A alone did not display AVP-induced currents **(Fig. 2D)**. In addition, AVP did not inhibit currents in oocytes expressing human SLO1/BK **(Supplementary Fig. 1).** Together, these findings demonstrate that activation of AVPR1A inhibits SLO2.1 channel activity.

**Figure 2.**
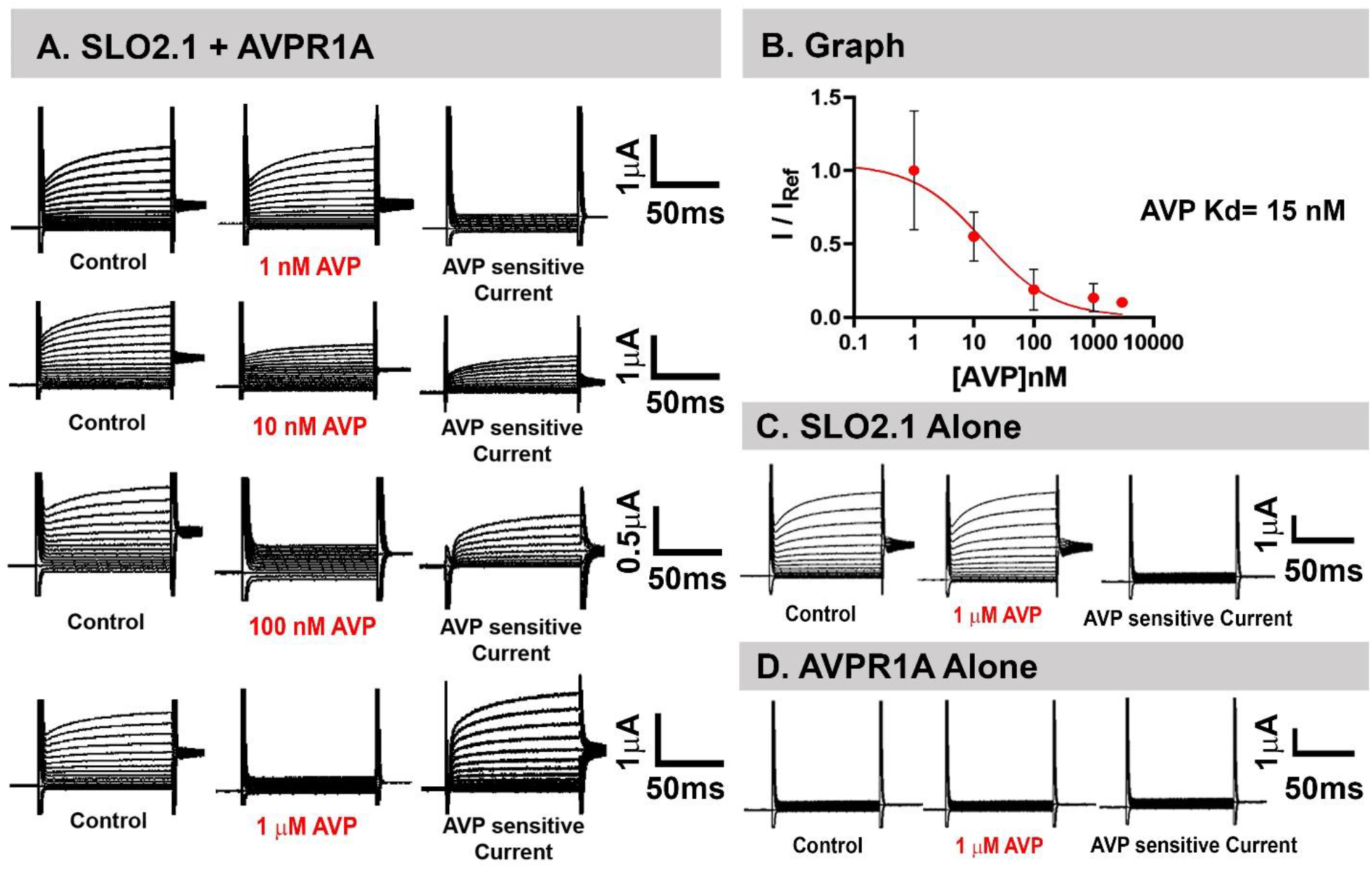
AVPR1A activation inhibits SLO2.1 currents in Xenopus oocytes. Whole-cell currents were measured in the two-electrode voltage clamp configuration. Oocytes were held at −80 mV, and currents were elicited by voltage steps from −100 to +100 mV. **A,** Representative current traces recorded from oocytes co-expressing human SLO2.1 and AVPR1A before and after application of AVP at 1, 10, 100, and 1,000 nM. **B,** Concentration-response relationship for AVP-dependent inhibition of SLO2.1-mediated currents in oocytes co-expressing SLO2.1 and AVPR1A. **C,** Representative currents from control oocytes expressing SLO2.1 without AVPR1A before and after AVP application. **D,** Representative recordings from oocytes expressing AVPR1A without SLO2.1 before and after AVP application.

### AVP reduces SLO2.1-dependent K⁺ currents in myometrial hTERT-HM cells

We next examined whether AVP modulates endogenous SLO2.1 currents in the immortalized human myometrial cell line hTERT-HM. Whole-cell and cell-attached patch-clamp recordings revealed outward K⁺ currents under basal conditions, and AVP exposure reduced these currents **(Fig. 3A,B)**. To determine whether the AVP-sensitive current was mediated by SLO2.1, we knocked down SLO2.1 expression with siRNA. SLO2.1 knockdown reduced the basal outward current and attenuated the inhibitory effect of AVP significantly more than in scrambled siRNA-treated cells **(Fig. 3C,D)**. These findings demonstrate that AVP inhibits a predominantly SLO2.1-mediated K⁺ current in hTERT-HM cells.

**Figure 3.**
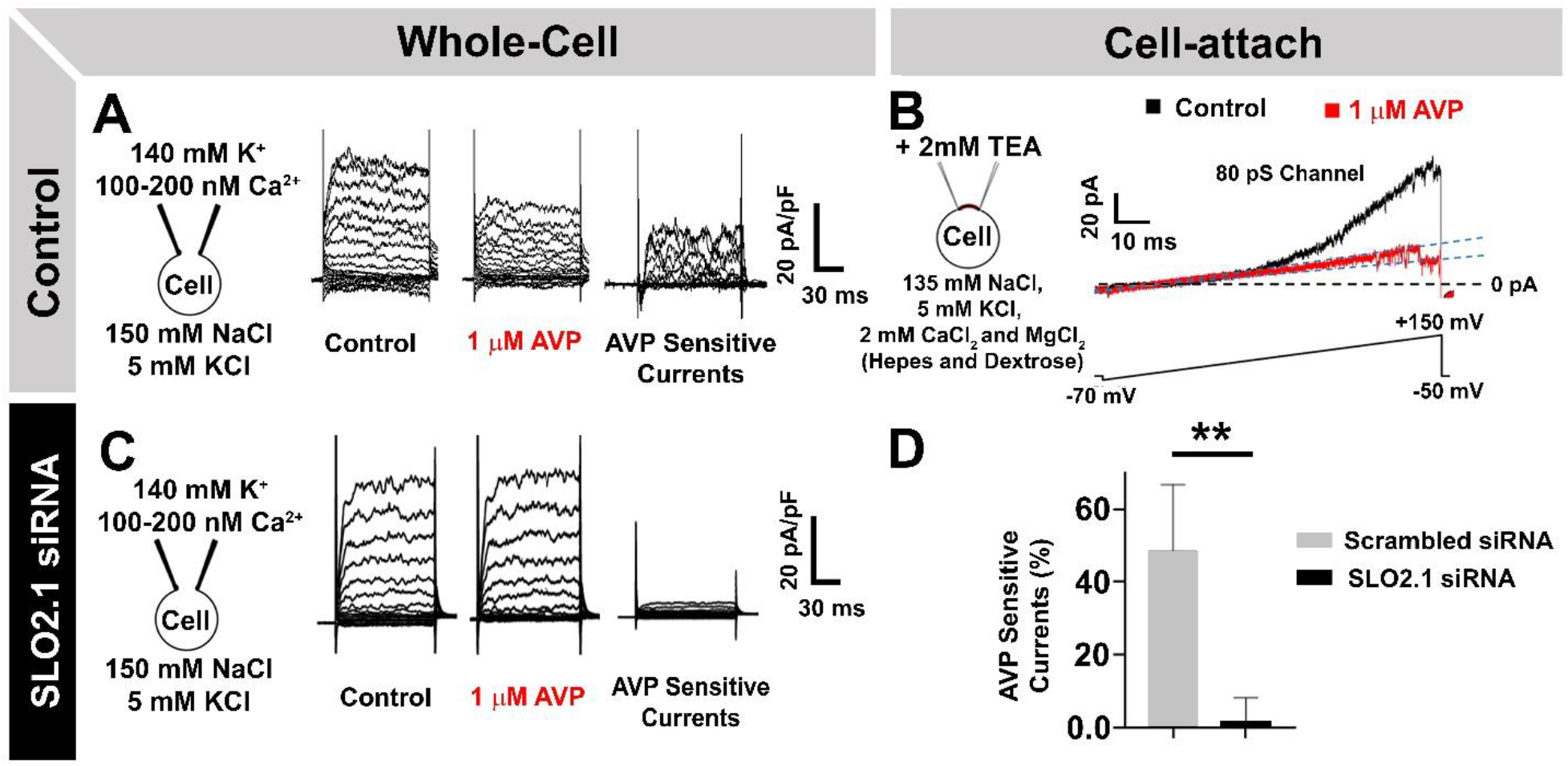
AVP inhibits SLO2.1-dependent K⁺ currents in hTERT-HM cells. **A,** Representative whole-cell K⁺ current traces recorded from hTERT-HM cells before and after AVP application. Cells were held at −70 mV, and currents were elicited by voltage steps from −90 to +150 mV. **B,** Representative cell-attached recordings obtained before and after AVP application. **C,** Representative whole-cell current traces from hTERT-HM cells treated with SLO2.1-targeting siRNA before and after AVP application. **D,** Quantification of AVP-sensitive whole-cell K⁺ currents in cells treated with scrambled siRNA or SLO2.1-targeting siRNA. AVP inhibited the current by 48.6 ± 18.1% in scrambled siRNA-treated cells (*n* = 4) and by 1.65 ± 1.89% in SLO2.1 siRNA-treated cells (*n* = 3). Data are presented as mean ± SD. \*\**P* < 0.01 by unpaired *t* test.

### AVP depolarizes hTERT-HM cells through AVPR1A- and SLO2.1-dependent mechanisms

Because inhibition of an outward K⁺ conductance is expected to depolarize the membrane potential, we measured membrane potential in hTERT-HM cells by using the voltage-sensitive dye DiSC₃(5) and flow cytometry. Responses were normalized to the hyperpolarization induced by the K⁺ ionophore valinomycin, which provided a consistent reference across experiments and treatment conditions **(Fig. 4A)**.

**Figure 4.**
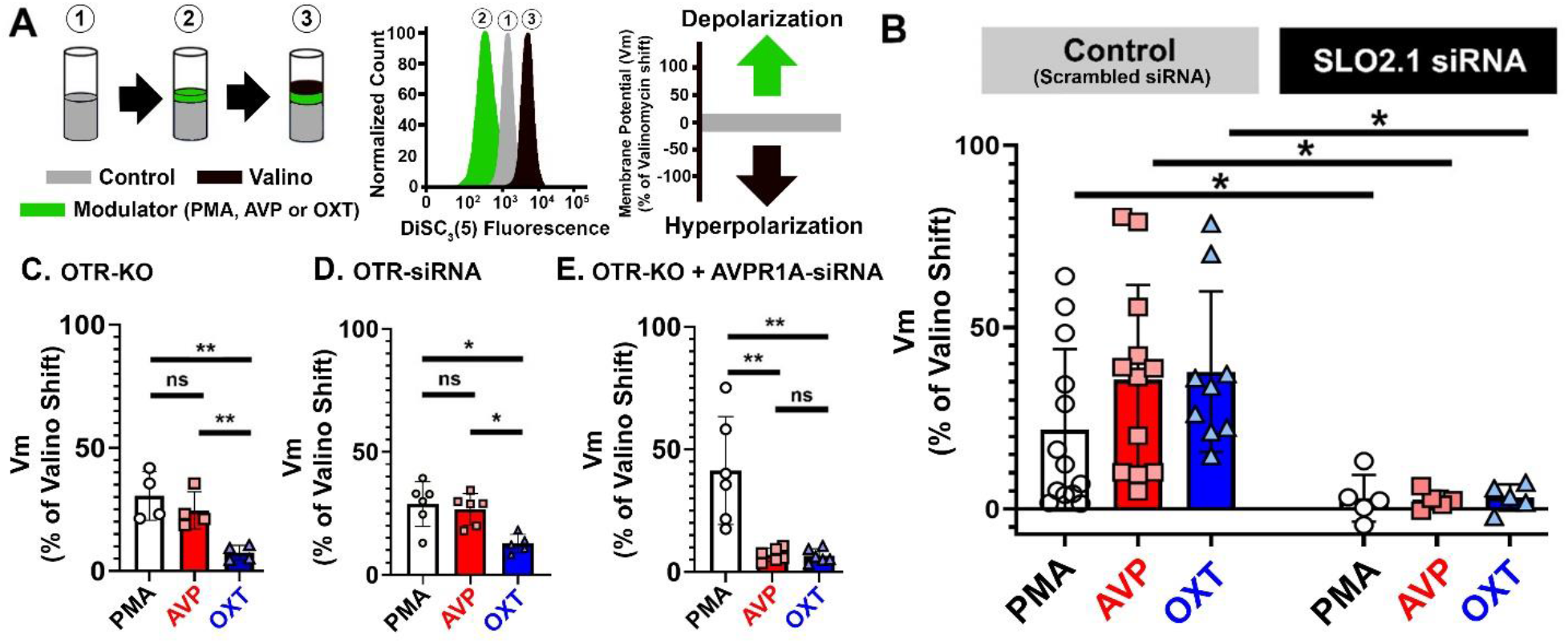
AVP depolarizes human myometrial hTERT-HM cells through AVPR1A- and SLO2.1-dependent mechanisms. Membrane potential (Vₘ) was assessed using the voltage-sensitive dye DiSC₃(5) and flow cytometry. **A,** Schematic of the experimental set-up, fluorescence response, and change in membrane potential (Vₘ). **B,** Effects of PMA, OXT, and AVP on Vₘ in control cells and cells treated with SLO2.1-targeting siRNA. Under control conditions, PMA, OXT, and AVP induced depolarizations of 21.79 ± 22.20%, 37.76 ± 22.10%, and 35.50 ± 26.10%, respectively. The AVP response was not significantly different from the OXT response (P = 0.833) or the PMA response (P = 0.173). After SLO2.1 knockdown, PMA-, OXT-, and AVP-induced depolarizations were reduced to 2.95 ± 6.40%, 3.19 ± 3.60%, and 2.49 ± 2.50%, respectively (n = 5 per treatment). **C,** Effects of PMA, OXT, and AVP in stable OTR-knockout cells. PMA, OXT, and AVP induced depolarizations of 30.50 ± 9.88%, 7.44 ± 2.99%, and 24.65 ± 7.56%, respectively (n = 4). **D,** Effects of PMA, OXT, and AVP in hTERT-HM cells treated with OTR-targeting siRNA. PMA, OXT, and AVP induced depolarizations of 28.79 ± 9.09%, 12.73 ± 3.76%, and 26.64 ± 6.36%, respectively (n = 6). **E,** Effects of PMA, OXT, and AVP in OTR-knockout cells treated with AVPR1A-targeting siRNA. PMA, OXT, and AVP induced depolarizations of 41.37 ± 21.99%, 6.62 ± 1.14%, and 6.98 ± 1.10%, respectively (n = 6). Data are presented as mean ± SD. \**P* < 0.05, \*\**P* < 0.01 by unpaired t tests; ns, not significant.

We first compared the effect of AVP with those of OXT and the protein kinase C activator phorbol 12-myristate 13-acetate (PMA), both of which inhibit SLO2.1 and depolarize hTERT-HM cells (Ferreira *et al*., 2019, 2021, 2025). PMA, OXT, and AVP each depolarized control cells, with comparable responses among the three treatments **(Fig. 4B left side)**. Knockdown of SLO2.1 markedly reduced the depolarization induced by all three treatments, indicating that AVP-induced and OXT-induced depolarization depends largely on inhibition of the SLO2.1-mediated hyperpolarizing current **(Fig. 4B right side)**.

To determine whether AVP-induced depolarization involved cross-activation of the oxytocin receptor (OTR), we examined responses in a stable OTR-knockout cell line and in hTERT-HM cells treated with OTR-targeting siRNA. OTR loss substantially reduced OXT-induced depolarization, whereas AVP- and PMA-induced depolarization were largely preserved **(Fig. 4 C,D)**. These findings indicate that the AVP response does not require OTR activation.

We next investigated whether AVPR1A mediated AVP-induced depolarization by treating OTR-knockout cells with AVPR1A-targeting siRNA. Combined disruption of OTR and AVPR1A markedly reduced the responses to both OXT and AVP, whereas the response to PMA remained intact **(Fig. 4E)**. Together, these findings demonstrate that AVP depolarizes hTERT-HM cells predominantly through AVPR1A and supports a mechanism in which AVPR1A-dependent PKC activation inhibits SLO2.1 channels, thereby promoting membrane depolarization.

### AVP induces biphasic Ca²⁺ signals in hTERT-HM cells

To determine whether AVP increases intracellular Ca²⁺ through mechanisms similar to those activated by OXT, we performed Fluo-4 AM Ca²⁺ imaging in hTERT-HM cells stimulated with 1 μM AVP or 1 μM OXT. In the presence of 2 mM extracellular Ca²⁺, both AVP and OXT induced a rapid initial Ca²⁺ peak followed by an oscillatory phase **(Fig. 5A,E)**. In nominally Ca²⁺-free solution, the initial Ca²⁺ peaks induced by both agonists were preserved, whereas the oscillatory phases were markedly reduced **(Fig. 5B,F,I,L)**. These findings indicate that the initial Ca²⁺ peak does not require extracellular Ca²⁺ entry, consistent with release from intracellular stores, whereas the oscillatory phase depends strongly on extracellular Ca²⁺ influx.

**Figure 5.**
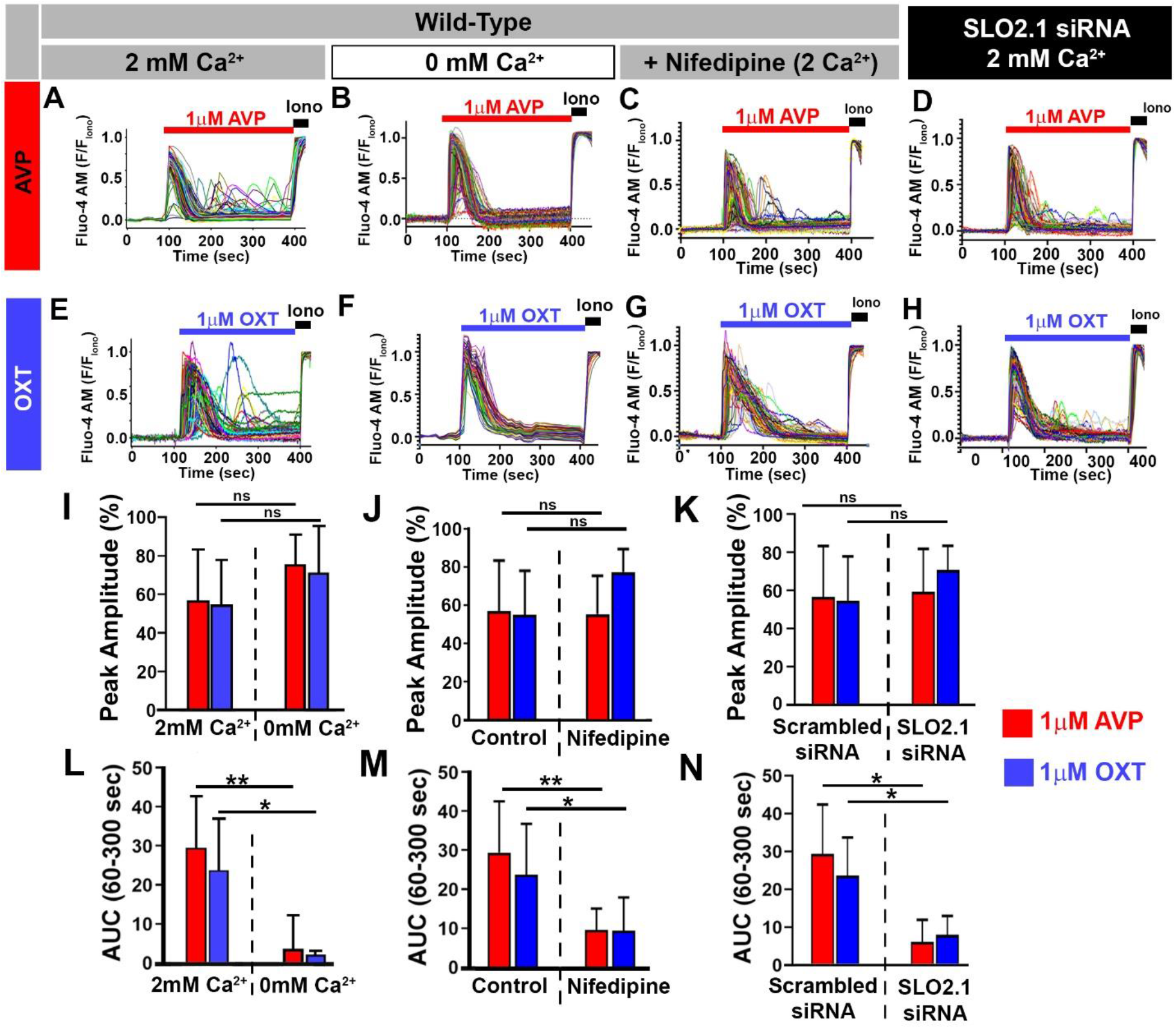
Effects of extracellular Ca²⁺, nifedipine, and SLO2.1 knockdown on AVP- and OXT-induced Ca²⁺ signals in hTERT-HM cells. Intracellular Ca²⁺ responses were measured using Fluo-4 AM imaging. **A,** Representative Ca²⁺ traces from cells stimulated with 1 μM AVP in the presence of 2 mM extracellular Ca²⁺ or **B,** nominally Ca²⁺-free external solution containing no added Ca²⁺ **C,** Representative Ca²⁺ trace from cells stimulated with 1 μM AVP in the presence of 2 mM extracellular Ca²⁺ and 10 μM nifedipine. **D,** Representative Ca²⁺ trace from cells treated with SLO2.1-targeting siRNA and stimulated with 1 μM AVP in the presence of 2 mM extracellular Ca²⁺**. E,** Representative Ca²⁺ traces from cells stimulated with 1 μM OXT in the presence of 2 mM extracellular Ca²⁺ or **F,** nominally Ca²⁺-free external solution containing no added Ca²⁺**. G,** Representative Ca²⁺ trace from cells stimulated with 1 μM OXT in the presence of 2 mM extracellular Ca²⁺ and 10 μM nifedipine. **H,** Representative Ca²⁺ trace from cells treated with SLO2.1-targeting siRNA and stimulated with 1 μM OXT in the presence of 2 mM extracellular Ca²⁺. **A–H,** Fluorescence values were normalized to the maximal fluorescence obtained with 5 μM ionomycin in the presence of 2 mM extracellular Ca²⁺**. I,** Quantification of the initial Ca²⁺ peak amplitude induced by AVP and OXT in the presence of 2 mM or 0 mM added extracellular Ca²⁺. In 2 mM extracellular Ca²⁺, peak amplitudes were 56.6 ± 26.7% for AVP and 54.6 ± 23.3% for OXT. In nominally Ca²⁺-free solution, peak amplitudes were 71.5 ± 23.9% for AVP and 75.7 ± 15.2% for OXT. **J,** Quantification of the initial Ca²⁺ peak amplitude induced by AVP and OXT in the absence or presence of 10 μM nifedipine. **K,** Quantification of the initial Ca²⁺ peak amplitude induced by AVP and OXT in cells treated with scrambled or SLO2.1-targeting siRNA. In scrambled siRNA-treated cells, peak amplitudes were 56.6 ± 26.7% for AVP and 54.6 ± 23.3% for OXT. Following SLO2.1 knockdown, peak amplitudes were 59.4 ± 22.4% for AVP and 70.8 ± 12.5% for OXT. **L,** Quantification of the oscillatory-phase area under the curve (AUC) in the presence of 2 mM or 0 mM added extracellular Ca²⁺. In 2 mM extracellular Ca²⁺, AUC values were 29.51 ± 13.07 for AVP and 23.82 ± 10.02 for OXT. In nominally Ca²⁺-free solution, AUC values were 3.98 ± 8.51 for AVP and 2.26 ± 1.04 for OXT. For the 2 mM extracellular Ca²⁺ control conditions, n = 7. **M,** Quantification of the oscillatory-phase AUC in the absence or presence of nifedipine. Nifedipine reduced the AUC from 29.51 to 9.78 for AVP and from 23.82 to 9.60 for OXT. For nifedipine-treated conditions, n = 5. **N,** Quantification of the oscillatory-phase AUC in cells treated with scrambled or SLO2.1-targeting siRNA. In scrambled siRNA-treated cells, AUC values were 29.51 ± 13.07 for AVP and 23.82 ± 10.02 for OXT. Following SLO2.1 knockdown, AUC values were 6.29 ± 5.79 for AVP and 8.13 ± 5.02 for OXT. For siRNA experiments, n = 5. Data are presented as mean ± SD. *P < 0.05, **P < 0.01 by unpaired t tests; ns, not significant.

To determine whether voltage-dependent Ca²⁺ channels contribute to either phase, cells were treated with nifedipine. Nifedipine had little effect on the initial Ca²⁺ peaks induced by AVP or OXT but significantly reduced the oscillatory-phase area under the curve (AUC) for both agonists **(Fig. 5C,G,J,M)**. These results support a major contribution of voltage-dependent Ca²⁺ entry to the oscillatory phase but not to the initial peak.

We next tested whether SLO2.1 contributes to either phase of the Ca²⁺ response. SLO2.1 knockdown did not alter the percentage of cells responding to AVP or OXT and did not reduce the initial Ca²⁺ peak induced by either agonist **(Fig. 5D,H,K)**. In contrast, SLO2.1 knockdown markedly reduced the oscillatory-phase AUC induced by both AVP and OXT **(Fig. 5N)**.

Together, these findings show that AVP and OXT induce biphasic Ca²⁺ responses in hTERT-HM cells. The initial phase is independent of SLO2.1 and extracellular Ca²⁺ entry and is consistent with release from intracellular stores. The subsequent oscillatory phase depends on SLO2.1-mediated membrane excitability and extracellular Ca²⁺ influx through voltage-dependent Ca²⁺ channels.

### AVP and OXT differentially regulate biphasic Ca²⁺ signaling

Given the biphasic Ca²⁺ responses described in **Fig. 5**, we investigated whether AVP and OXT differ in their concentration-dependent activation of the initial Ca²⁺ peak and the subsequent oscillatory phase. hTERT-HM cells were stimulated with AVP or OXT at concentrations ranging from 0.01 nM to 1 μM in the presence of 2 mM extracellular Ca²⁺. Responses were quantified according to three parameters: the percentage of responding cells, the amplitude of the initial Ca²⁺ peak, and the AUC of the oscillatory phase measured from 60 to 360 s after stimulation. OXT elicited Ca²⁺ responses at lower concentrations than AVP. The EC₅₀ for the percentage of responding cells was 0.16 nM for OXT and 40.47 nM for AVP, whereas the maximal percentages of responding cells were similar for the two agonists, reaching 87.53% for OXT and 89.45% for AVP **(Fig. 6A,B).** OXT was also more potent than AVP in increasing the amplitude of the initial Ca²⁺ peak. The EC₅₀ values for peak amplitude were 0.110 nM for OXT and 0.745 nM for AVP, with maximal normalized peak amplitudes of 0.550 and 0.594, respectively **(Fig. 6C).** In contrast, AVP was more potent than OXT in promoting the oscillatory phase. The EC₅₀ for the oscillatory-phase AUC was 0.0667 nM for AVP and 0.422nM for OXT, whereas the maximal AUC values were comparable at 25.80 for AVP and 23.47 for OXT **(Fig. 6D).** Thus, OXT was more potent at triggering the initial Ca²⁺ peak, whereas AVP more potently promoted the later oscillatory component associated with extracellular Ca²⁺ entry.

**Figure 6.**
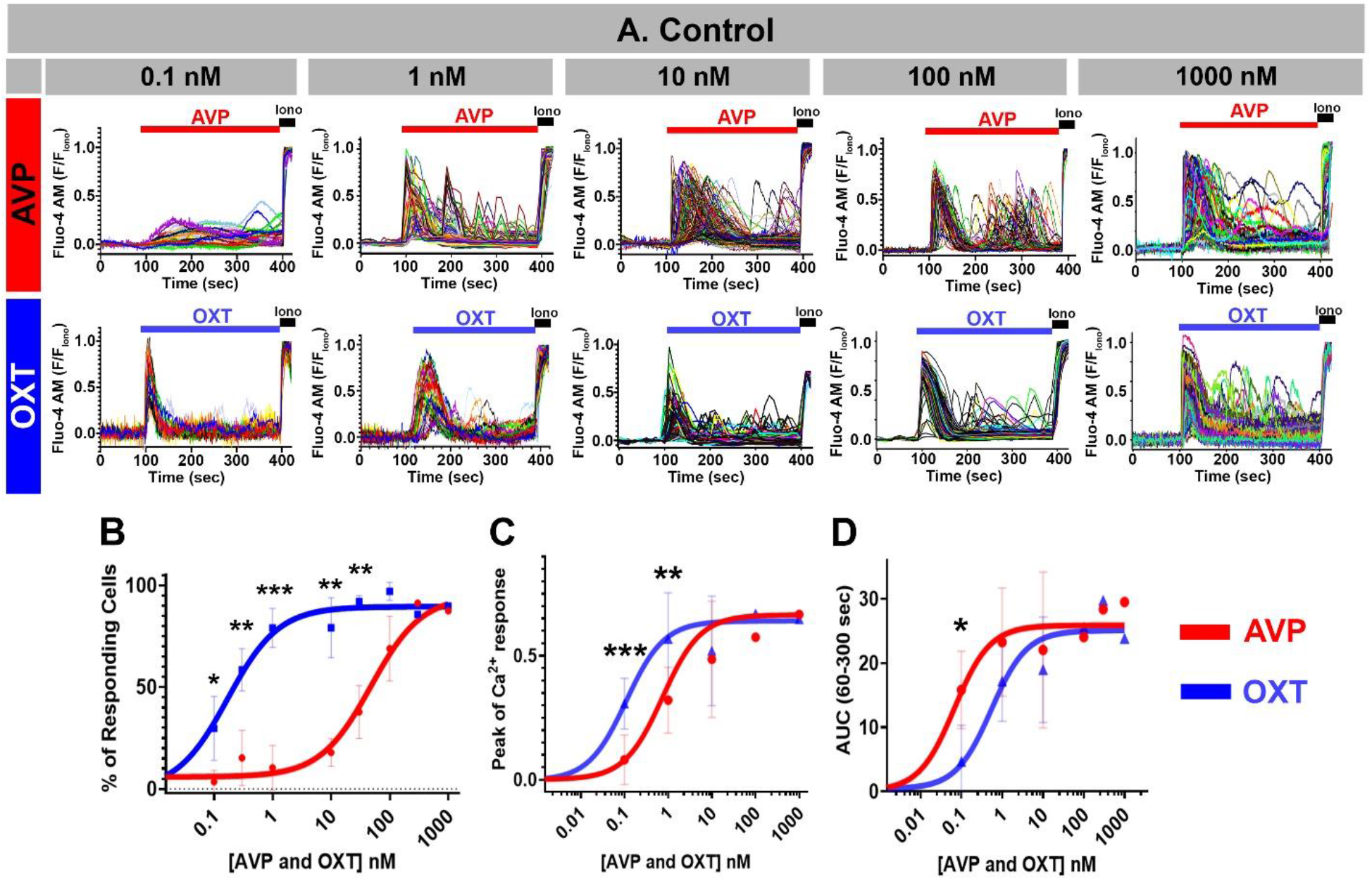
Concentration-dependent intracellular Ca²⁺ responses to AVP and OXT in hTERT-HM cells. Intracellular Ca²⁺ responses were measured with Fluo-4 AM imaging in the presence of 2 mM extracellular Ca²⁺. **A,** Representative Ca²⁺ traces from hTERT-HM cells stimulated with AVP or OXT at concentrations ranging from 0.1 pM to 1 μM. Fluorescence values were normalized to the maximal fluorescence obtained with 5 μM ionomycin in the presence of 2 mM extracellular Ca²⁺. **B,** Concentration-response relationships for the percentage of responding cells. The fitted EC₅₀ values were 40.47 nM for AVP and 0.1607 nM for OXT, with maximal responses of 89.45% and 87.53%, respectively. **C,** Concentration-response relationships for the initial Ca²⁺ peak amplitude. The fitted EC₅₀ values were 0.745 nM for AVP and 0.110 nM for OXT, with maximal normalized peak amplitudes of 0.594 and 0.550, respectively. **D,** Concentration-response relationships for the area under the curve (AUC) of the oscillatory phase measured from 60 to 360 s after stimulation. The fitted EC₅₀ values were 0.0667 nM for AVP and 0.4216 nM for OXT, with maximal AUC values of 25.80 and 23.47, respectively. Concentration-response curves were fitted by nonlinear regression using the three-parameter agonist-versus-response model in GraphPad Prism. For AVP, the number of independent experiments was *n* = 15 at 0.1 pM; *n* = 5 at 0.1, 0.3, 1, 10, 30, 100, and 300 nM; and *n* = 7 at 1 μM. For OXT, the number of independent experiments was *n* = 12 at 0.1 pM; *n* = 4 at 0.1, 0.3, 1, 10, and 30 nM; *n* = 3 at 100 and 300 nM; and *n* = 7 at 1 μM. Data are presented as mean ± SD. \**P* < 0.05, \*\**P* < 0.01, \*\*\**P* < 0.001 by One-Way Anova.

## Discussion

Together, our findings support a model in which AVP-mediated activation of AVPR1A in myometrial cells stimulates both IP₃-dependent Ca²⁺ release from intracellular stores and voltage dependent calcium influx through PKC-dependent inhibition of SLO2.1. Inhibition of SLO2.1 reduces outward K⁺ conductance, depolarizes the membrane potential, and promotes extracellular Ca²⁺ entry through voltage-dependent Ca²⁺ channels **(Fig. 7)**. Several observations support this model. First, SLO2.1, OTR, AVPR1A, and AVPR2 were detected in primary MSMCs and in hTERT-HM cells. AVPR2 was less abundant in term labor samples than in term non-labor samples, suggesting that AVP signaling during labor is primarily through AVPR1A. Second, AVPR1A activation was sufficient to inhibit SLO2.1 channel activity in *Xenopus* oocytes, whereas AVP had no direct effect on SLO2.1 currents in the absence of AVPR1A. Third, in human myometrial cells, AVP reduced outward K⁺ currents, and this effect was markedly attenuated after SLO2.1 knockdown. Fourth, AVP depolarized the resting membrane potential of MSMCs, and this response was reduced after knockdown of either SLO2.1 or AVPR1A, supporting the conclusion that AVPR1A-dependent inhibition of SLO2.1 contributes to AVP-induced membrane depolarization. Fifth, AVP and OXT induced biphasic Ca²⁺ responses consisting of an initial Ca²⁺ peak followed by a later oscillatory phase. The oscillatory phase was reduced by removal of extracellular Ca²⁺, by nifedipine, and by SLO2.1 knockdown, indicating that this later calcium increase depends on membrane depolarization and extracellular Ca²⁺ entry. Finally, OXT was more potent in initiating the early Ca²⁺ response, whereas AVP more effectively promoted the oscillatory phase, suggesting differential regulation of the initial store-release phase and the sustained Ca²⁺ entry-dependent phase. Although the EC₅₀ for AVP-induced Ca²⁺ oscillations was lower than the IC₅₀ for SLO2.1 inhibition measured in Xenopus oocytes, these values were obtained in different experimental systems and are therefore not directly comparable.

**Figure 7.**
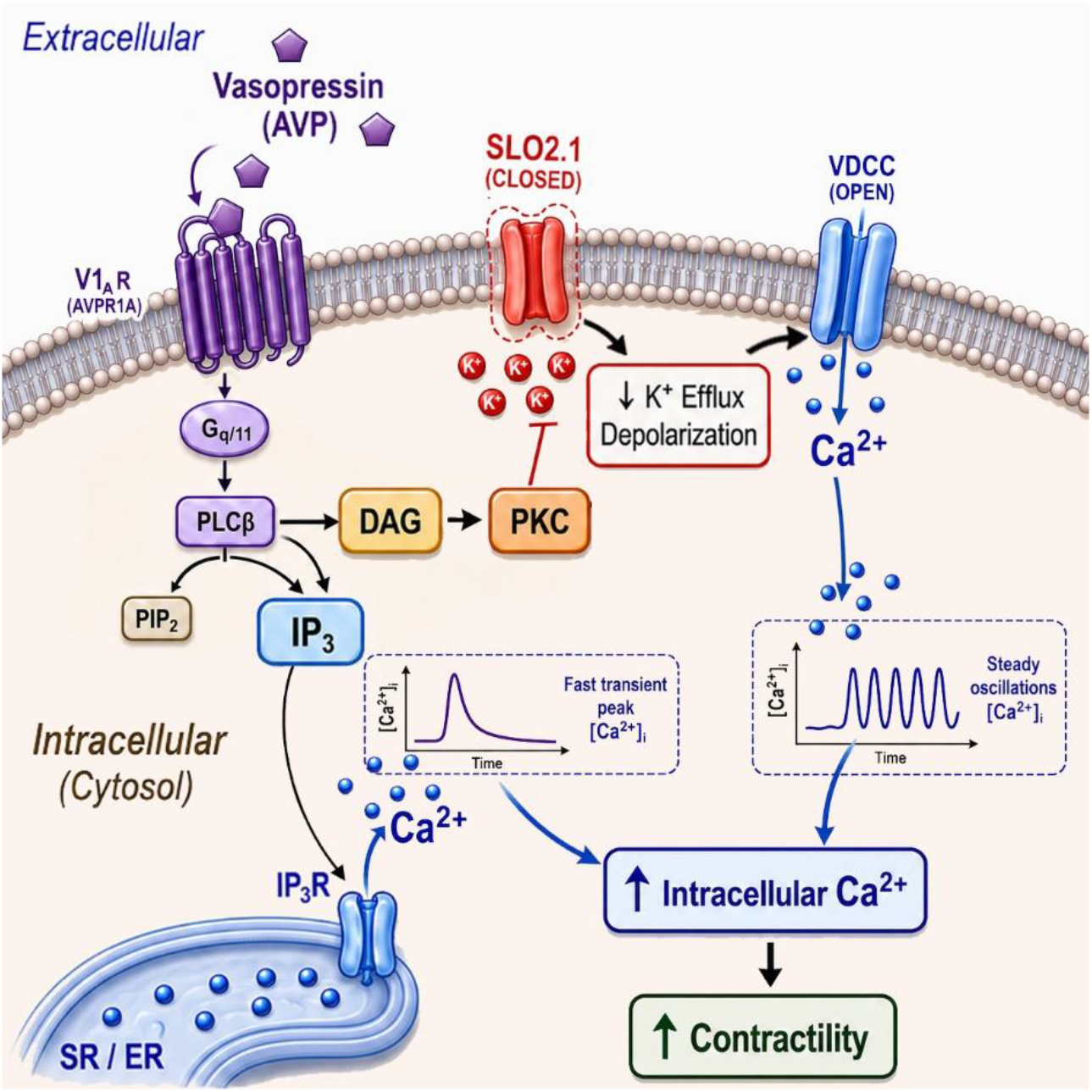
Proposed model of AVP-dependent uterotonic signaling in human myometrial smooth muscle cells. AVP binds to AVPR1A, a Gq/11-coupled receptor, activating phospholipase C (PLC) and promoting generation of inositol 1,4,5-trisphosphate (IP₃) and diacylglycerol (DAG). IP₃ activates IP₃ receptors (IP3R) on sarcoplasmic reticulum/endoplasmic reticulum (SR/ER), producing an initial release of Ca²⁺ into the cytosol. In parallel, DAG activates protein kinase C (PKC), which inhibits K⁺ conductance through SLO2.1. Suppression of SLO2.1 reduces the hyperpolarizing K⁺ outward current that normally stabilizes the resting membrane potential in a quiescent state and thereby promotes membrane depolarization. This depolarization increases the activity of voltage-dependent Ca²⁺ channels (VDCCs), promoting extracellular Ca²⁺ entry and overall cell contractility.

Although our experiments were designed to define the cellular mechanism of AVP action rather than its role in specific pregnancy disorders, the pathway identified here may be relevant under conditions associated with elevated or sustained AVP signaling. Elevated AVP concentrations have been linked to pregnancy complications such as preeclampsia (Santillan *et al*., 2014; Scroggins *et al*., 2018). AVP release can also be stimulated by pain, stress, hypoxia, and bleeding, conditions that may occur during complicated pregnancies (KENDLER, WEITZMAN and FISHER, 1978; Suzuki *et al*., 2009; Sivukhina *et al*., 2010; Szczepanska-Sadowska *et al*., 2017). Although excessive uterine contractions are not a primary feature of preeclampsia (Pitaphrom and Sukcharoen, 2006; TIKKANEN, 2011; Ananth *et al*., 2016), abnormal uterine activation, including tachysystole (Heuser *et al*., 2013), may be clinically significant in this context because excessive contractions can further reduce placental blood flow and increase maternal and fetal risk. Future studies examining AVPR1A half-life, trafficking, and receptor recycling in myometrial smooth muscle cells will be important for understanding how sustained or repeated AVP signaling may influence the risk of these complications.

Outside of pregnancy, the AVPR1A–SLO2.1 pathway may be relevant to primary dysmenorrhea, a condition characterized by excessive uterine activity and altered AVP signaling. Women with primary dysmenorrhea exhibit elevated circulating AVP concentrations, increased myometrial activity, and reduced uterine blood flow. In the non-pregnant uterus, AVP is a particularly potent uterotonic stimulus, and uterine sensitivity and AVPR1A receptor density increase during the premenstrual phase (Åkerlund, 2002). ReaMoreover, experimentally increasing circulating AVP enhances uterine contractility and pain in women with dysmenorrhea, whereas antagonists that block AVPR1A and OTR, including atosiban and SR 49059, have shown therapeutic effects (Åkerlund, 2002). Our findings provide a potential ion-channel mechanism for these observations. AVPR1A-dependent inhibition of SLO2.1 would reduce outward K⁺ conductance, promote membrane depolarization, and enhance voltage-dependent Ca²⁺ entry, thereby favoring sustained myometrial excitability and contraction. In combination with AVP-mediated vasoconstriction and reduced uterine perfusion, this pathway could contribute to the uterine hyperactivity, ischemia, and pain associated with primary dysmenorrhea. Further studies are needed to determine whether SLO2.1 activity or AVPR1A–SLO2.1 coupling is altered across the menstrual cycle or in women with dysmenorrhea.

Interpretation of these physiological and pathological implications also requires consideration of receptor selectivity, because OXT and AVP and their corresponding receptors exhibit substantial structural and pharmacological overlap (Postina, Kojro and Fahrenholz, 1996; Chini and Manning, 2007; Stoop, 2012; Rae *et al*., 2022; Soumier *et al*., 2022). OTR and AVPR1A share substantial sequence similarity, particularly within the transmembrane domains and extracellular loops that contribute to ligand binding (Tribollet and Barberis, 1996; Hawtin *et al*., 2005; Wootten *et al*., 2011, 2018). Conserved residues within these regions are critical for OXT and AVP binding, supporting the possibility that each hormone may interact with both receptor families under specific conditions. Consistent with this, *in vitro* studies have shown that AVP can bind and activate both vasopressin and oxytocin receptors. OXT can also interact with vasopressin receptors, although generally with lower affinity than AVP (MAGGI *et al*., 1990; Maggi *et al*., 1991; Busnelli *et al*., 2013).

Despite this potential for receptor crosstalk, our findings indicate that AVP responses in hTERT-HM cells are largely independent of OTR activation, at least under the experimental conditions used here. Specifically, both cells with full OTR knockout and cells with siRNA-mediated OTR knockdown showed a marked reduction in OXT-induced membrane potential changes, but no significant reduction in AVP-induced responses. Similarly, in hTERT-HM OTR knockout cells, OXT-induced calcium signals were strongly reduced, whereas AVP-induced calcium responses were largely preserved. These findings support the interpretation that, in this experimental system, AVP primarily signals through AVPR1A.

In summary, our findings support a model in which AVP increases myometrial excitability through AVPR1A-dependent inhibition of SLO2.1. This work extends the concept that uterotonic GPCRs regulate myometrial function not only through canonical IP₃-mediated Ca²⁺ release from intracellular stores, but also by modulating ion channels that control membrane potential and sustained Ca²⁺ entry. By identifying SLO2.1 as a downstream effector of AVPR1A signaling, our study highlights a mechanism by which AVP may contribute to abnormal uterine activation under pathological conditions. From a therapeutic perspective, preserving SLO2.1 activity or limiting AVPR1A-dependent inhibition of SLO2.1 may be a strategy to stabilize uterine excitability and reduce excessive myometrial activation.

## Acknowledgments

We thank Dr. Deborah Frank, scientific editor in the Department of Obstetrics and Gynecology, for critical review of the manuscript. We also thank the Division of Clinical Research staff in the Department of Obstetrics and Gynecology at Washington Universitiy School of Medicine for obtaining patients’ consent and acquiring human myometrial biopsies. This work was supported by National Institutes of Health grant R01HD088097 (to C.M.S. and S.K.E.), to and the Department of Obstetrics and Gynecology at Washington University in St. Louis.

