## Supplementary Figures for "Activation of Vasopressin Receptor 1A by Vasopressin Enhances Myometrial Smooth Muscle Cell Excitability by Inhibiting the Potassium Channel SLO2.1"

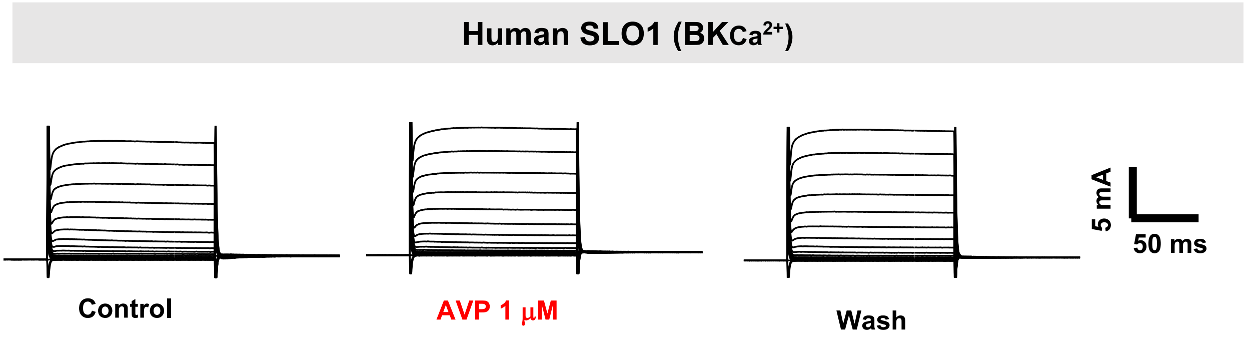


**Supplementary Figure 1. AVP does not induce $K^+$ currents in the absence of AVP receptor.** Representative two-electrode voltage-clamp recordings from Xenopus oocytes expressing human SLO1 before and after application of 1 μM AVP.

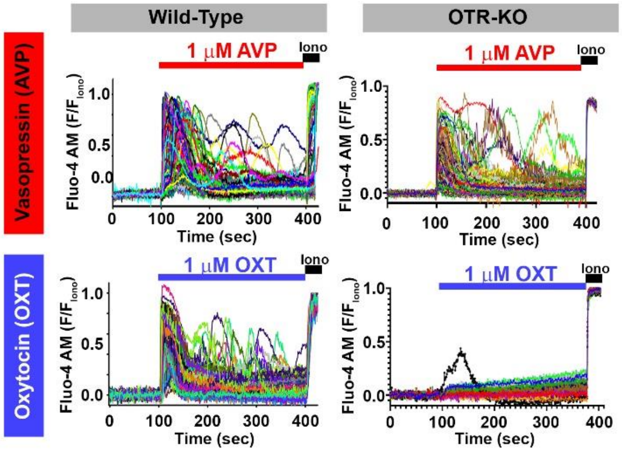


**Supplementary Figure 2. AVP and OXT induce $Ca^{2+}$ signals in hTERT-HM Wild type cells and in hTERT OTR KO cells.** Representative Fluo-4 AM $Ca^{2+}$ imaging traces from hTERT-HM Wild Type cells **(Top left)** or from Oxytocin receptor KO hTERT-HM Cells (OTR-KO) **(Top right),** both stimulated with 1 μM AVP in the presence of 2 mM extracellular $Ca^{2+}$**.** Representative Fluo-4 AM $Ca^{2+}$ imaging traces from hTERT-HM Wild Type cells **(Bottom left)** or from OTR-KO hTERT-HM Cells **(Bottom right),** both stimulated with 1 μM OXT in the presence of 2 mM extracellular $Ca^{2+}$**.** All values were normalized to fluorescence obtained with 5 μM ionomycin (Iono) in 2 mM extracellular $Ca^{2+}$.
